# Colchicine Inhibits Smooth Muscle Cell Phenotypic Switch and Aortic Dissection in Mice

**DOI:** 10.1101/2024.11.02.621511

**Authors:** Hui Jiang, Yaping Zhao, Mei Jin, Zhidan Zhang, Li Wang, Zhiping Liu, Suowen Xu, Jianping Weng

## Abstract

Aortic dissection is a serious cardiovascular disease characterized by intimal tearing and vascular delamination. Smooth muscle cell phenotypic transformation, extracellular matrix degradation and vascular inflammation are the main pathogenic mechanisms of aortic dissection. Colchicine is an alkaloid derived from the plant Colchicum autumnale L. and was the first FDA-approved anti-inflammatory drug in the cardiovascular field. In this study, we investigated the protective effects of colchicine in aortic dissection in mice and found that colchicine resisted aortic dissection by inhibiting vascular inflammation and smooth muscle cell phenotypic transformation, and that colchicine reversed smooth muscle cell phenotypic transformation by targeting the transcriptional activator myocardin. This study provides new directions for the development of drug therapies for aortic dissection.

## Introduction

Aortic dissection is a cardiovascular condition where a tear in the intimal layer allows blood to separate the vessel wall layers, causing delamination. Open and endovascular repair surgery is the primary treatment option^1^. The pathogenesis involves vascular smooth muscle cells (VSMCs) transitioning from a contractile to a synthetic phenotype, along with inflammatory cell infiltration that releases matrix metalloproteinases and cytokines, promoting extracellular matrix degradation and dissection progression^2^. Colchicine is an alkaloid compound found in the *Colchicum* plant, recognized for its potent anti-inflammatory effects and considered for treating various cardiovascular diseases, such as pericarditis, atrial fibrillation, and coronary syndromes^3^. The FDA has approved colchicine as the first anti-inflammatory drug for the treatment of atherosclerotic cardiovascular disease. However, it remains unknown whether colchicine can inhibit the development of aortic dissection.

## Methods

To investigate the protective effect of colchicine in aortic dissection, we used a β -Aminopropionitrile (BAPN)-induced aortic dissection model in mice. Three-week-old male C57BL/6J mice were fed a standard chow diet, and BAPN (A3134, Sigma-Aldrich, USA) was administered via drinking water at a concentration of 1 g/kg/day for 3 weeks. Simultaneously, mice were gavaged with either colchicine (200 μg/kg/day) (C3915, Sigma-Aldrich, USA) or an equivalent dose of saline (Fig. A). All mouse experiments were approved by Institutional Animal Care and Use Committee. The antibodies used in the experiments including α-SMA (sc-56499, Santa Cruz), SM22-α (GTX628831, GeneTex), CNN1 (13938-1-AP, Thermo Fisher), TUBULIN (66031-1-Ig, Proteintech), GAPDH (60004-1-lg, Proteintech).

## Results

### Colchicine inhibits aortic dissection in mice

Mice were monitored daily for survival, and any mice died were immediately dissected to examine the aorta. Mice in the colchicine group showed a higher survival rate. Gross evaluation of the aorta revealed that the control group exhibited a severe disease phenotype with an 80% incidence of dissection, whereas the colchicine-treated group showed a significantly lower incidence of dissection at 36% (Fig. A). Pathological analysis revealed that the vehicle group developed significant dissections characterized by intimal tearing and the formation of a false lumen. H&E staining showed erythrocytes passing through the lumen, filling the detached vascular layers, while EVG staining indicated elastic lamina fragmentation. In contrast, the aortic vessels in the colchicine group displayed improved structural integrity (Fig. B).

### RNA sequencing profiling revealed that colchicine inhibits vascular inflammation and smooth muscle cell phenotypic switch

To understand mechanistic insights into Colchicine-mediated protective effects, transcriptome sequencing analysis was performed in diseased aortic tissues from control and colchicine-treated mice (data are available at NCBI Gene Expression Omnibus (GEO): accession# GSE279545). Differentially expressed genes (DEGs) (log2(FC) > 1 and p < 0.05), identified 502 significantly upregulated genes and 1,471 significantly downregulated genes (Fig. C). Reactome pathway enrichment analysis of these DEGs revealed significant enrichment in pathways related to the immune system and signal transduction. Analysis of molecular genes related to smooth muscle cell switch and vascular inflammation showed that contractile phenotype-related genes (*Myocd, Myh11, Cnn1, Acta2*) were significantly upregulated in the colchicine-treated group, while synthetic phenotype-related genes (*Fn1, Spp1*), inflammation-related genes (*Il1b, Il6, Ccl2*), and extracellular matrix degradation genes (*Mmp9*) were markedly reduced compared to the vehicle group (Fig. C). These findings suggest that colchicine may exert its protective effects by reducing vascular inflammation and inhibiting phenotypic switch of smooth muscle cells.

**Figure.**
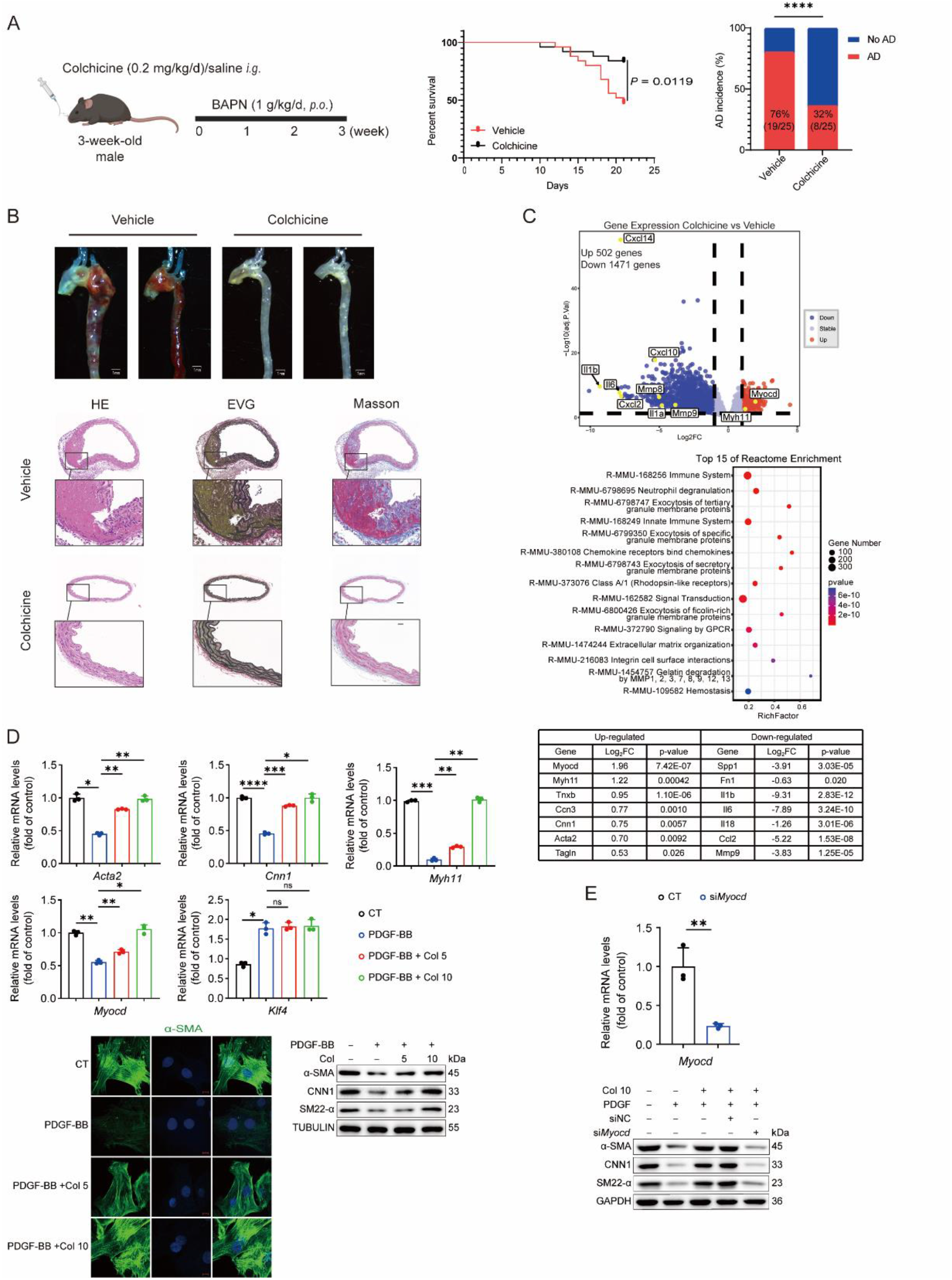
Colchicine resists aortic dissection in mice. (A) Schematic representation of aortic dissection induction in three-week-old male mice by administering β-aminopropionitrile monofumarate (BAPN) in drinking water, along with colchicine or saline gavage treatment; Survival curves of colchicine-treated and control mice. Survival rates were analyzed using the Log-rank (Mantel-Cox) test (n = 25 per group); Incidence of aortic dissection in colchicine-treated and control mice. Incidence was compared using Fisher’s exact test (n = 25 per group). (B) Representative images of the aortas from colchicine-treated and control mice (n = 25 per group); Representative histological sections of the aorta from colchicine-treated and control mice stained with hematoxylin-eosin (HE), elastic van Gieson (EVG), and Masson’s trichrome (Masson). Scale bar, 100 μm for original image; Scale bar, 20 μm for enlarged inset. (C) Volcano plot of differentially expressed genes in the aorta of colchicine-treated versus control mice. Blue dots indicate downregulated genes, while red dots represent upregulated genes; Reactome pathway enrichment analyses of differentially expressed genes in the aortas of colchicine-treated and control mice; Levels of smooth muscle cell phenotype-switching and inflammatory genes in colchicine-treated mice compared to control mice. (D) Relative mRNA levels of contractile markers and transcription factors in rat vascular smooth muscle cells under PDGF-BB stimulation and treated with various concentrations of colchicine (5 nM and 10 nM) (n = 3 per group). Statistical significance: *p < 0.05, **p < 0.01, ***p < 0.001; one-way ANOVA; In another set of experiments, immunofluorescence staining of α-SMA was performed in rat smooth muscle cells were treated with different concentrations of colchicine (5 nM and 10 nM) in the absence or presence of PDGF-BB (n = 4 per group); Scale bar, 10 μm; Western blot analysis of contractile protein markers levels in rat vascular smooth muscle cells stimulated by PDGF-BB and treated with different concentrations of colchicine (5 nM and 10 nM) (n = 3 per group). (E) Relative mRNA levels of *Myocd* in rat vascular smooth muscle cells under siRNA of *Myocd*; Western blot analysis of contractile markers in PDGF-BB stimulated rat vascular smooth muscle cells treated with 10 nM colchicine after *Myocd* knockdown (si*Myocd*) (n = 3 per group).

### Colchicine targets MYOCD to reverse smooth muscle cell phenotypic switch

To further investigate the mechanism by which colchicine inhibits the development of aortic dissection, we examined its ability to suppress platelet-derived growth factor BB (PDGF-BB)-induced phenotypic switch in primary rat VSMCs. Colchicine effectively reversed the PDGF-BB-induced downregulation of VSMC contractile molecules at both the gene level—including *Acta2, Cnn1, Myh11*, and *Myocd*—and the protein level, including α-SMA, CNN1, SM22-α (Fig. D). Since MYOCD is a co-activator of serum response factor, a transcription factor that regulates the contractile phenotype of VSMCs^4^, we hypothesized that colchicine may influence the expression of downstream VSMC contractile molecules by modulating MYOCD expression. To test this hypothesis, we employed MYOCD-specific siRNA. Upon knockdown of MYOCD, the ability of colchicine to reverse the VSMC phenotype was lost, suggesting that MYOCD is a key target of colchicine’s action (Fig. E). Additionally, colchicine did not exhibit a significant inhibitory effect on TNF-α-induced inflammation in VSMCs (data not shown).

## Discussion

In conclusion, this study highlights that colchicine, a well-known anti-inflammatory drug, holds significant promise for the treatment of aortic dissection, a critical cardiovascular disease. In a mouse model of aortic dissection, colchicine markedly improved survival rates by inhibiting the phenotypic switch of VSMCs and reducing vascular inflammation.

Since colchicine’s known mechanism involves binding to microtubule proteins and inhibiting their polymerization, which suppresses inflammatory cell recruitment and cytokine release^5^, further research is needed to confirm whether colchicine reduces inflammation by affecting immune cells in aortic dissection. Although there is a lack of clinical trials of colchicine for the treatment of aortic dissection, results of clinical trials have demonstrated that long-term low-dose colchicine (0.5 mg/day) has a favorable safety profile in the treatment of atherosclerosis with no significant side effects^6^. However, the clinically effective dose of colchicine for the treatment of aortic dissection is unknown and requires more diverse animal models and clinical trials to confirm its efficacy and the lowest effective dose in treating aortic dissection.

## Declarations

## Acknowledgments

S.X. is Senior Humboldt Research Fellow of Alexander von Humboldt Foundation, Germany. This study was supported by grants from the National Key R&D Program of China (Grant No. 2021YFC2500500), the National Natural Science Foundation of China (Grant Nos. 82370444, 82070464, 82270500, 12411530127). This work was also supported by the Program for Innovative Research Team of The First Affiliated Hospital of USTC (CXGG02) and Anhui Provincial Natural Science Foundation (Grant No. 2208085J08).

## Author contributions

Conceptualization, Suowen Xu, Jianping Weng, Zhiping Liu; Investigation, Hui Jiang, Mei Jin, Yaping Zhao; Writing – original draft, Yaping Zhao; Writing – review and editing, Zhidan Zhang, Suowen Xu, Li Wang, Zhiping Liu, Jianping Weng.

